# Modeling Stress-Induced Responses: Plasticity in Continuous State Space and Gradual Clonal Evolution

**DOI:** 10.1101/2023.07.03.547523

**Authors:** Anuraag Bukkuri

**Affiliations:** Cancer Biology and Evolution Program and Department of Integrated Mathematical Oncology, Moffitt Cancer Center

**Keywords:** Eco-Evolutionary Dynamics, Cancer Evolution, Bacterial Evolution, Plasticity, Therapeutic Resistance

## Abstract

Mathematical models of cancer and bacterial evolution have generally stemmed from a gene-centric framework, assuming clonal evolution via acquisition of resistance-conferring mutations and selection of their corresponding subpopulations. More recently, the role of phenotypic plasticity has been recognized and models accounting for phenotypic switching between discrete cell states (e.g. epithelial and mesenchymal) have been developed. However, seldom do models incorporate both plasticity and mutationally-driven resistance, particularly when the state space is continuous and resistance evolves in a continuous fashion. In this paper, we develop a framework to model plastic and mutational mechanisms of acquiring resistance in a continuous, gradual fashion. We use this framework to examine ways in which cancer and bacterial populations can respond to stress and consider implications for therapeutic strategies. Although we primarily discuss our framework in the context of cancer and bacteria, it applies broadly to any system capable of evolving via plasticity and genetic evolution.

## 1 Introduction

Therapeutic resistance is well-recognized as one of the major contributors to treatment failure and poor outcomes in patients [52, 37, 97, 108]. Traditionally, resistance is thought to emerge from the selection of subpopulations of cells that have acquired one or more resistance-conferring mutations [47]. However, recent studies also show the importance of phenotypic plasticity as a mechanism for mediating resistance, e.g., by cells entering a polyaneuploid, mesenchymal, inflammatory, or stem-like state [3, 18, 43, 55, 59, 71, 72, 101, 107, 109].

In order to develop novel therapeutic strategies for bacterial and cancer extinction or control, a deep understanding of their ecological (population), evolutionary (resistance), and demographic (state) dynamics is required. To this end, many mathematical models have been constructed and analyzed to elucidate various aspects of therapeutic resistance via plasticity and clonal evolution. However, most models consider only one of these mechanisms [9, 25, 33, 95]. For the few models that incorporate both [11, 12, 21], plastic transitions are usually assumed to occur within a discrete state space.

In this paper, we use integral projection models (IPMs) to develop a method to model the ecological, evolutionary, and demographic (hereafter referred to as eco-evo-demo) dynamics of populations that evolve resistance in the face of therapy via continuous phenotypic transitions and clonal evolution. IPMs developed as an off-shoot from matrix population models [10, 14, 15, 16, 24] as a way to deal with population structure of a continuous nature, relaxing the requirement that organisms in a population must be classified into a set of discrete states [28, 77]. These models allow ecologists and demographers to understand how continuous state structure impacts a population’s eco-evolutionary dynamics [19, 20]. IPMs are typically constructed from regression models that infer how an organism’s state and environment impacts its survival, growth, and reproduction [1, 22, 23, 64, 69]. Such models have been used in a variety of ecological contexts, to help us understand and control invasive species [29, 44], the adaptation of insects and plants to climate change [86], herbivore-plant interactions [50, 67, 79, 104], optimal flowering size in plants [51, 66, 68, 78, 103], and host-parasite dynamics [65]. We port over a more qualitative version of this framework to the cellular level and investigate the eco-evodemo dynamics in populations subjected to environmental stress. We also investigate two facultative mechanisms by which cell populations may modulate their stress-induced responses depending on their condition in the environment: stress-induced mutagenesis [4, 7, 34, 32] and facultative plasticity [2, 3, 31, 42, 60, 71, 72, 73, 74, 75, 89, 90]. Using this knowledge, we then examine the efficacy of intermittent and adaptive therapy strategies [40, 102].

## 2 Methods

### 2.1 Analytical Framework

We construct our modeling framework using IPMs. First, we outline the ecological dynamics for the most simple case in the form of an integrodifference equation:

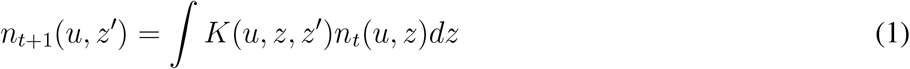

where *n* is the population distribution, *u* is the drug resistance trait, *z ∈ Z* = [0, 1] is the current state of the cell, and *z*^*′*^ is state at the next time step. The states *z ∈ Z* may represent the state of cells on epithelial to mesenchymal, adrenergic (ADRN) to mesenchymal (MES), or aneuploidy continua, for example. The kernel *K* can be decomposed into survival, plasticity, birth, and reproduction as follows:

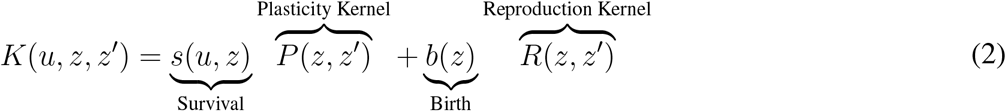

The first term is the transition kernel, often denoted by *P* (*u, z, z*^*′*^), and the second term is the fecundity kernel, *F* (*z*^*′*^, *z*). The survival and birth functions, *s*(*u, z*) and *b*(*z*), are called *individual components* whereas the plasticity and reproduction kernels, *P* (*z, z*^*′*^) and *R*(*z, z*^*′*^), are *state-redistribution components*. Under this formulation, we assume that drug resistance, *u*, only impacts the survival of the cells–in future sections, we will relax this assumption. The specific forms for each of these components are:

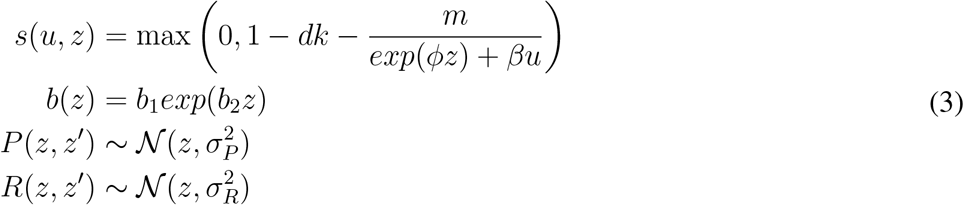

Critical to note is the presence of a survival-birth trade-off. Higher cell states (e.g., mesenchymal, inflammatory, or high ploidy) attain higher survival under therapy than lower cell states (e.g., epithelial, adrenergic, proliferative, or low ploidy). But this comes at a cost: a lower birth rate. Such a trade-off is experimentally observed in a multitude of contexts across bacterial and cancer evolution [3, 18, 55, 60, 59, 85, 91, 109]. From these ecological dynamics, we can then derive the evolutionary dynamics of resistance via Fisher’s fundamental theorem of natural selection [58, 5, 36, 57], *sensu* the *G* function framework [9, 98]. This approach states that evolution is the product of evolvability (the capacity to generate heritable variation upon which natural selection can act), and the selection gradient. Namely, for the discrete-time case, we have:

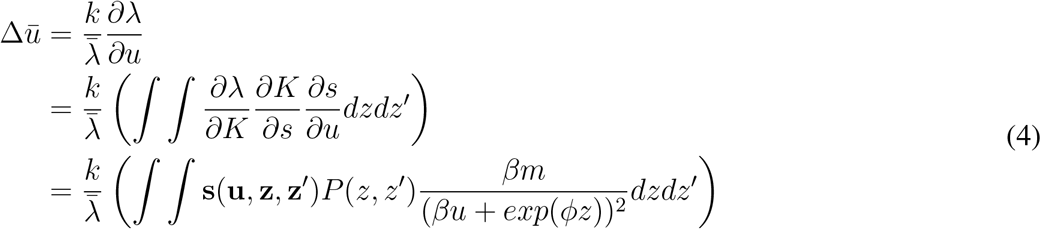

where 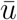 represents the mean drug resistance value in the population, *λ* is the per-capita growth rate or fitness function calculated as the spectral radius of the kernel operator [10], and **s**(**u, z, z**^*′*^) is the sensitivity of the fitness function to perturbations in the kernel operator, calculated as the normalized inner product of the reproductive and stable state distribution eigenvectors. Note that if drug resistance levels impact components other than just survival, this equation will need to be modified to incorporate the other terms via a basic application of the chain rule as shown above. Under this framework, we allow cells to respond to therapeutic stressors via both plasticity (changing cell state in a continuous and reversible fashion) and genetic evolution (evolving higher levels of resistance, *u*).

### 2.2 Numerical Implementation

There are two technical points to note concerning the numerical implementation of our model. The discretized IPM kernel used in the simulations is the result of overlaying a mesh over the operator, treating it as a large, finite-dimensional matrix. The choice of mesh size (and thereby dimensionality of the resulting matrix) is a trade-off between accuracy and computational cost: The smaller the mesh size, the more accurate the numerical results, but the greater computational effort. In our case, we used a 100x100 matrix. Another aspect to consider are the boundary conditions. Since we defined a bounded interval for our state distribution, *Z*, we must correct for eviction, whereby individuals diffuse outside these boundaries due to the Gaussian nature of our size-redistribution kernels. To do this, we return all cells that occupy states outside *Z* to the boundary from which they were evicted [64] (see [105] for alternative corrections). Another consequence of our modeling assumptions is the existence of a small subpopulation of highly resistant cells occupying high cell states prior to drug exposure. Due to the Gaussian nature of our kernels, these cells can indeed become or generate drug-sensitive cells and vice versa. This observation has found experimental support, e.g., Shaffer et al. identify a rare population of melanoma cells that transiently display high expression of AXL in the absence of therapy. These cells are resistant to anti-BRAF therapy and have been found to stochastically give rise to drug-sensitive cells [87].

## 3 Results

We ran a series of simulations capturing the eco-evo-demo dynamics of populations under therapy. In addition, supplementary movies for each plot are provided to visualize how the distribution and densities of cell states in the population change over time. The parameter values used in the simulations can be found in Table 1. These parameters were chosen to be biologically plausible, numerically convenient for simulation purposes, and to clearly show differences among stress-responses and therapeutic outcomes. Although many of these parameters can quantitatively alter results (e.g., a higher birth rate and evolvability or a lower drug dosage and natural death rate can promote the survival of the cell population under therapy), the same broad qualitative trends hold.

**Table 1:**
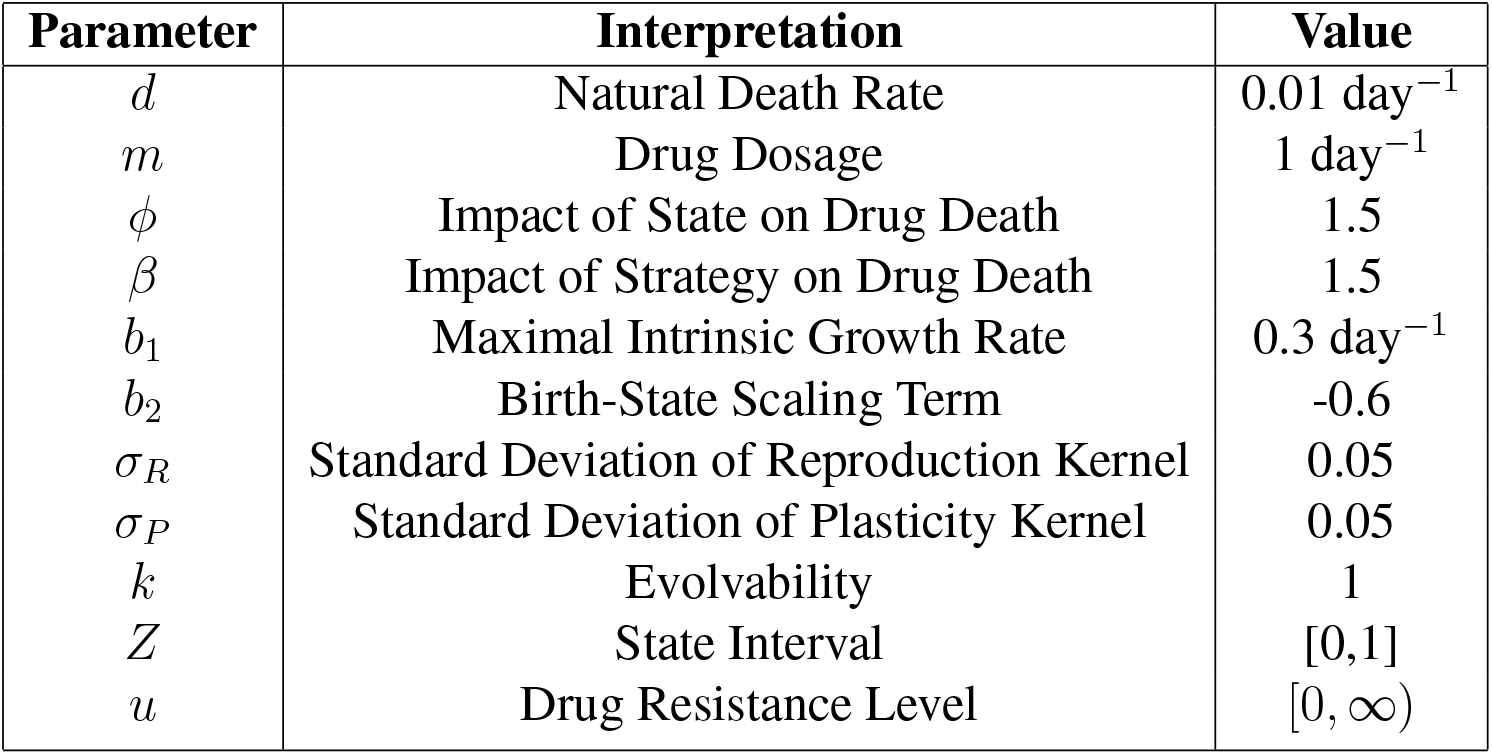
Parameter definitions and baseline values used in simulations.

| Parameter | Interpretation | Value |
| --- | --- | --- |
| $d$ | Natural Death Rate | $0.01 \text{ day}^{-1}$ |
| $m$ | Drug Dosage | $1 \text{ day}^{-1}$ |
| $\phi$ | Impact of State on Drug Death | 1.5 |
| $\beta$ | Impact of Strategy on Drug Death | 1.5 |
| $b_1$ | Maximal Intrinsic Growth Rate | $0.3 \text{ day}^{-1}$ |
| $b_2$ | Birth-State Scaling Term | -0.6 |
| $\sigma_R$ | Standard Deviation of Reproduction Kernel | 0.05 |
| $\sigma_P$ | Standard Deviation of Plasticity Kernel | 0.05 |
| $k$ | Evolvability | 1 |
| $Z$ | State Interval | [0,1] |
| $u$ | Drug Resistance Level | $[0, \infty)$ |

In all simulations, the population is initialized with densities of 0.5 across all cell states for a total population size of 50 cells. Therapy will be as a constant dose through time once the population reaches one million cells and the simulation will be stopped once the population recovers back to the one million cell threshold or once the population size is less than 10 (an arbitrary threshold). Key summary statistics are provided in Table 2: final drug resistance level, minimum of population, time of therapeutic application, time to progression (TTP: number of time steps between the start of therapy and when the population reaches one million cells again), and time to treatment failure (TTF: number of time steps between the start of therapy and when the population starts increasing in size).

**Table 2:** Summary statistics: model exploration.

|  | Baseline | Low Variance | High Variance | No Evolvability | High Evolvability |
| --- | --- | --- | --- | --- | --- |
| Drug Resistance | 3.850 | 4.068 | 3.556 | 0 | 4.868 |
| Minimum of Population | 38443 | 24375 | 68137 | 9 | 203847 |
| Start of Therapy | 48 | 48 | 48 | 46 | 53 |
| Time to Progression | 84 | 90 | 74 | NA | 33 |
| Time to Treatment Failure | 29 | 32 | 25 | NA | 10 |

### 3.1 Control Simulations

First, we run a simulation to understand the ecological and demographic dynamics of the population under therapy assuming no evolution of resistance by setting *k* = 0 (Fig. 1). Since cells in a lower state proliferate more rapidly than those in a higher state, the average cell state decreases from 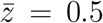 to 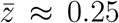 before therapy is administered. After 46 time steps, the population size passes the one million cell threshold and therapy is administered. In response, the population’s demography shifts towards higher cell states (Supplementary Movie 1), allowing the cells to minimize the effects of therapy at the cost of proliferating more slowly. This trade-off between death due to drug treatment and proliferation rate results in a leveling off of the demographic dynamics near 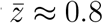. But a shift in the population’s demography is not enough for the population to avoid extinction.

**Figure 1:**
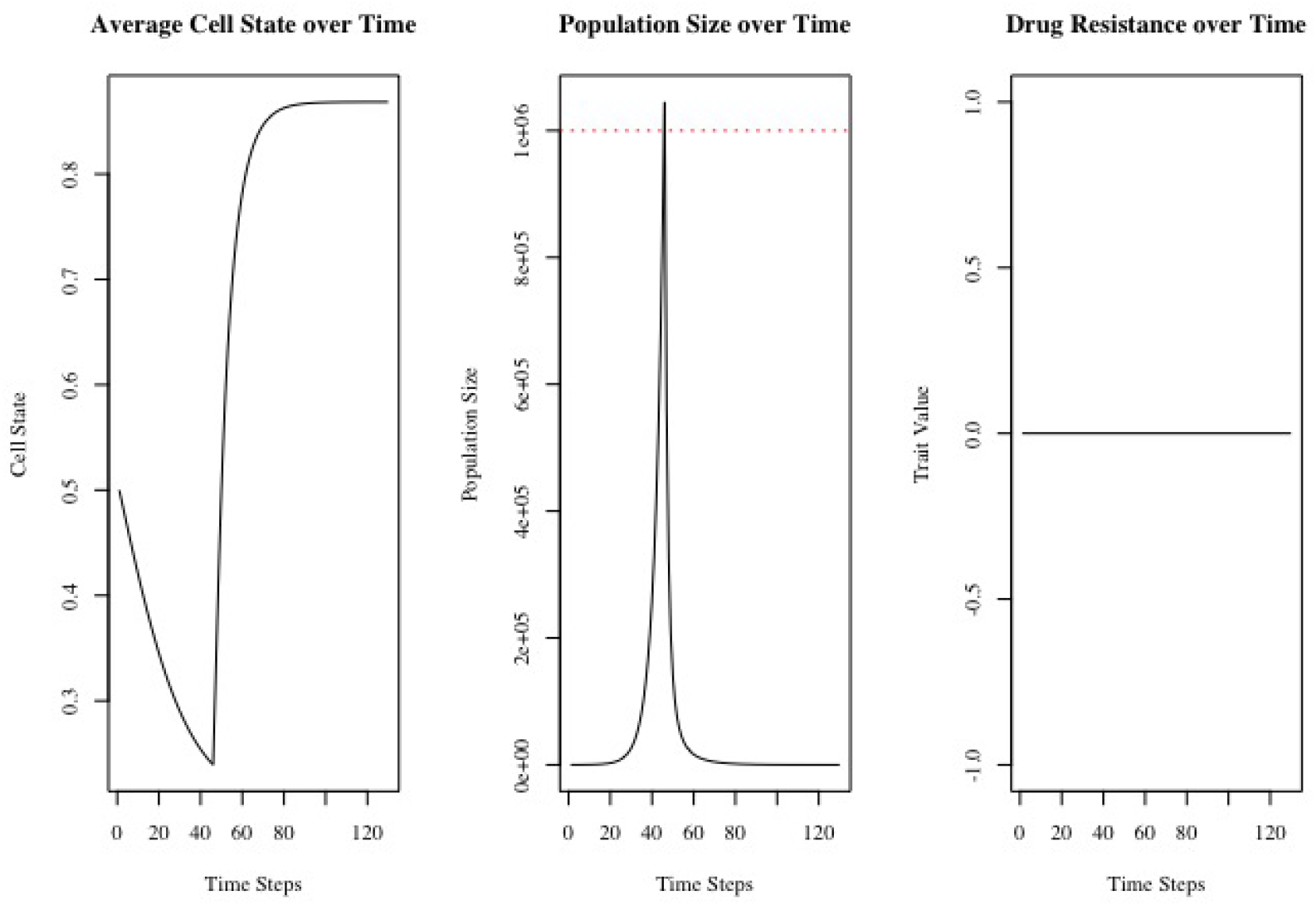
Eco-evo-demo dynamics without the evolution of resistance. Average cell state decreases before therapy is administered and increases after. Demographic dynamics begin to equilibrate due to the trade-off between escaping the effects of therapy and a low proliferation rate.

Next, we allow for the evolution of resistance by setting *k* = 1 and assess the resulting changes to the eco-evo-demo dynamics (Fig 2). As before, the average cell state decreases drastically before therapy is added due to the difference in birth rates for cells in low and high cell states. Due to the cost of evolvability, it takes two more time steps for the population to exceed one million cells. Once therapy is given, the population size drops dramatically and the population shifts its demography towards higher cell states (Supplementary Movie 2). This buys time for adaptive genetic evolution to occur [26, 35, 63]. Note that although adaptive plasticity aids in population persistence, it also slows down the rate of evolution by decreasing the selection pressure on the population [17]. As resistance develops and treatment begins to fail, the population shifts towards pre-therapy demography, leading to higher proliferation rates and the eventual progression of the tumor (Table 2).

**Figure 2:**
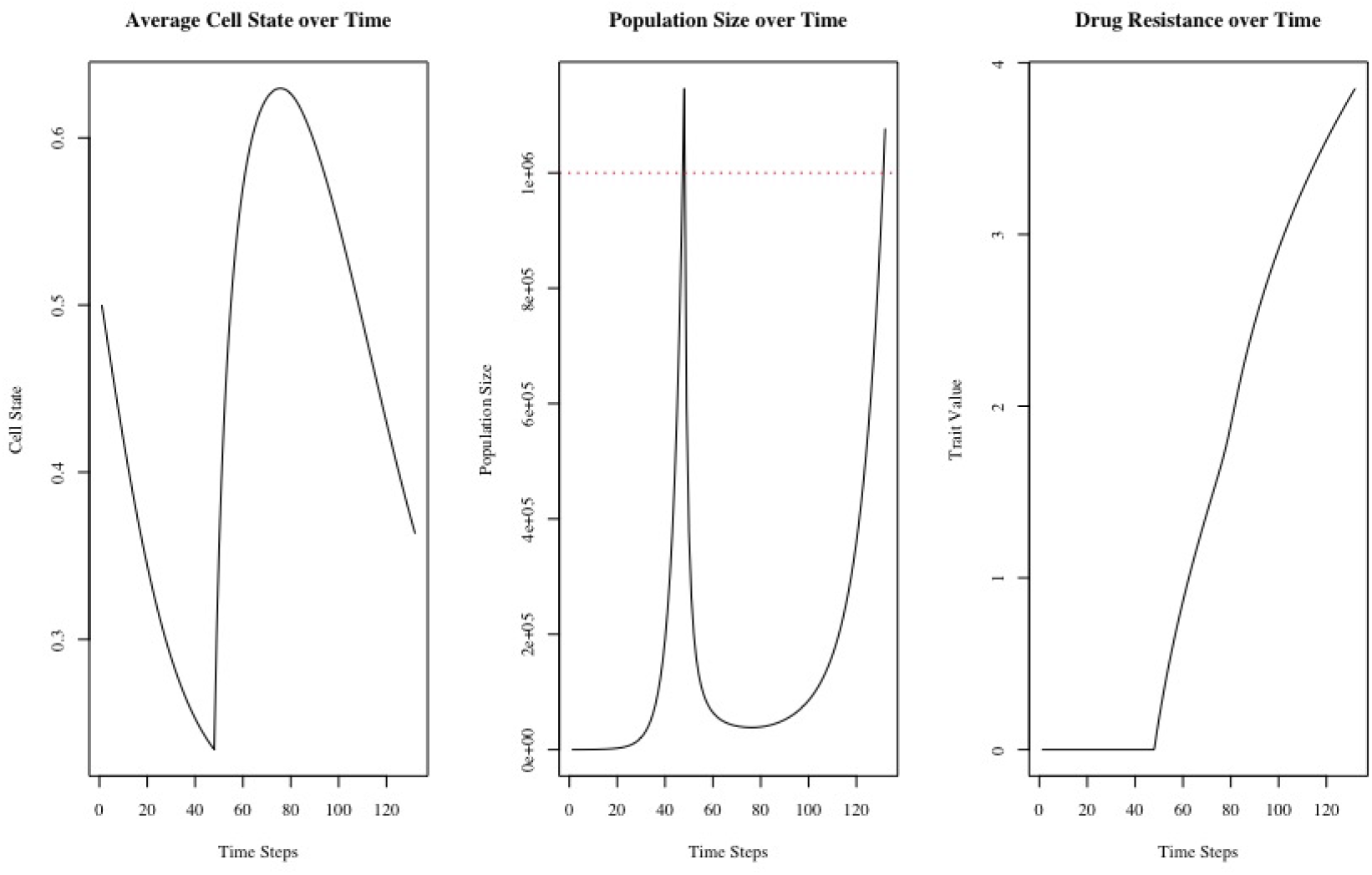
Eco-evo-demo dynamics with the evolution of resistance. Average cell state decreases before therapy is applied and increases immediately after. As resistance evolves, the demographic dynamics shift cells back towards lower cell states.

### 3.2 Model Exploration

Now, we run simulations to understand how variance and evolvability impact eco-evolutionary dynamics of populations under therapy. We start with the former. For biological realism and simplicity, we fix the variance in the fecundity kernel and solely consider changes to the plasticity kernel. An increase in the variance of the fecundity kernel can be considered a form of saltational evolution [106], *sensu* Goldschmidt’s “hopeful monster” macromutations [41] (a rather controversial theory [27], particularly at the cellular level). Combining this with the fact that the fecundity kernel produces variance across states, not the resistance trait, we choose to focus solely on the impacts of variance in the plasticity kernel. However, due to the structure of our model, results for fecundity variance are qualitatively similar to those of plasticity variance, with quantitative differences arising result from the magnitude difference between survival and birth probabilities and the interplay with evolutionary dynamics (survival is dependent on strategy but birth is not).

To assess the impact of variance in the plasticity kernel, we run simulations for low (*σ*_*G*_ = 0.01) and high (*σ*_*G*_ = 0.1) plasticity variance (Fig. 3A,B). Lower variance kernels can move closer to the tails of their distributions more effectively than higher variance kernels, a phenomenon that is paralleled in the pre-therapy demographic dynamics (Supplementary Movies 3,4). As expected, the average cell state in the population decreases until therapy is administered. However, the low (high) plasticity variance population reaches a lower (higher) average cell state than the baseline population (Fig 2). Due to the discrete-time nature of the simulation, the low, baseline, and high plasticity variance populations all begin treatment at the same time. However, lower plasticity variance populations grow faster and reach a higher population size upon the application of therapy.

**Figure 3:**
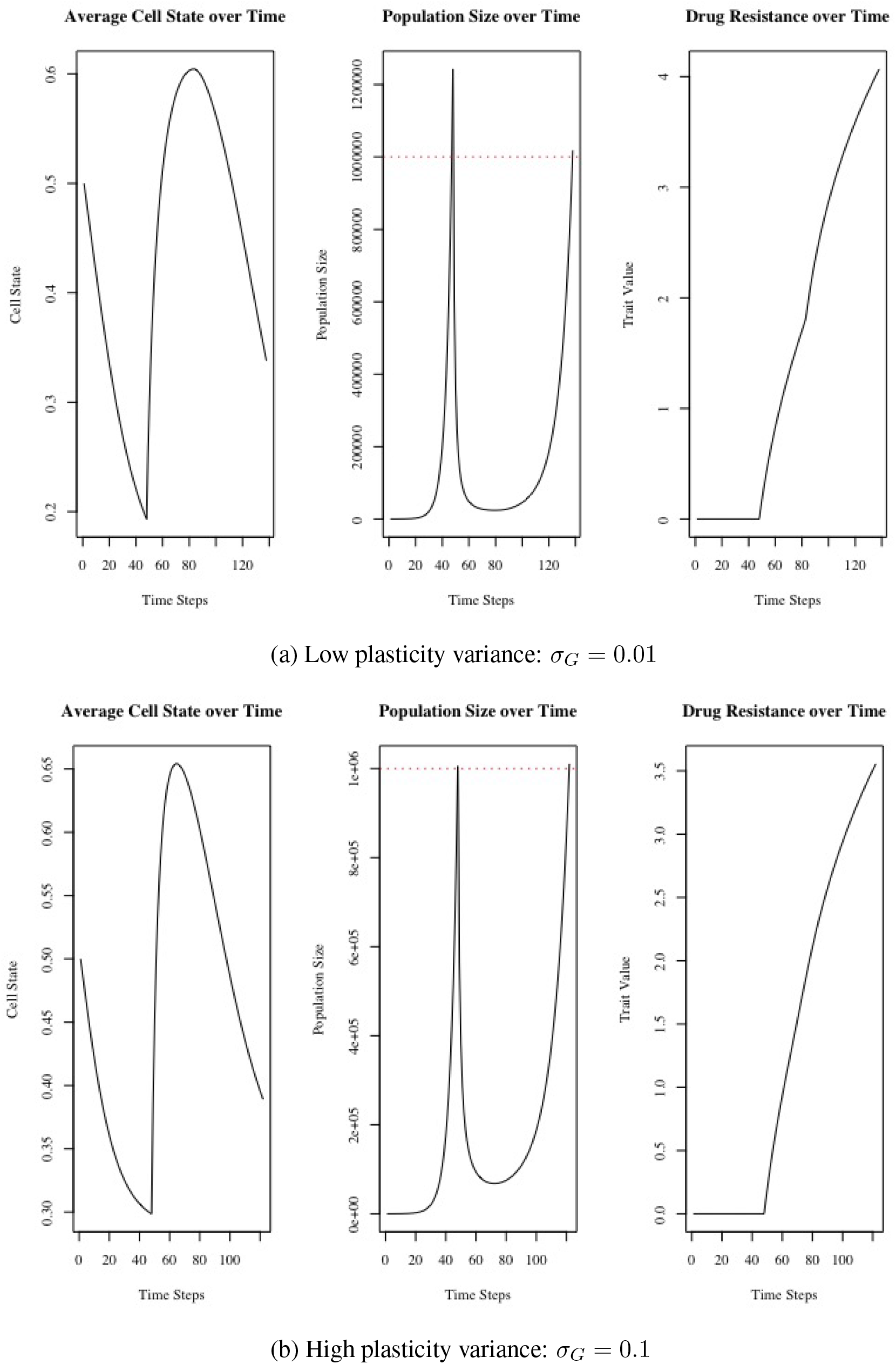
Impact of plasticity variance on eco-evo-demo dynamics. Higher plasticity variances lead to higher population minima and faster times to progression and treatment failure.

**Figure 4:**
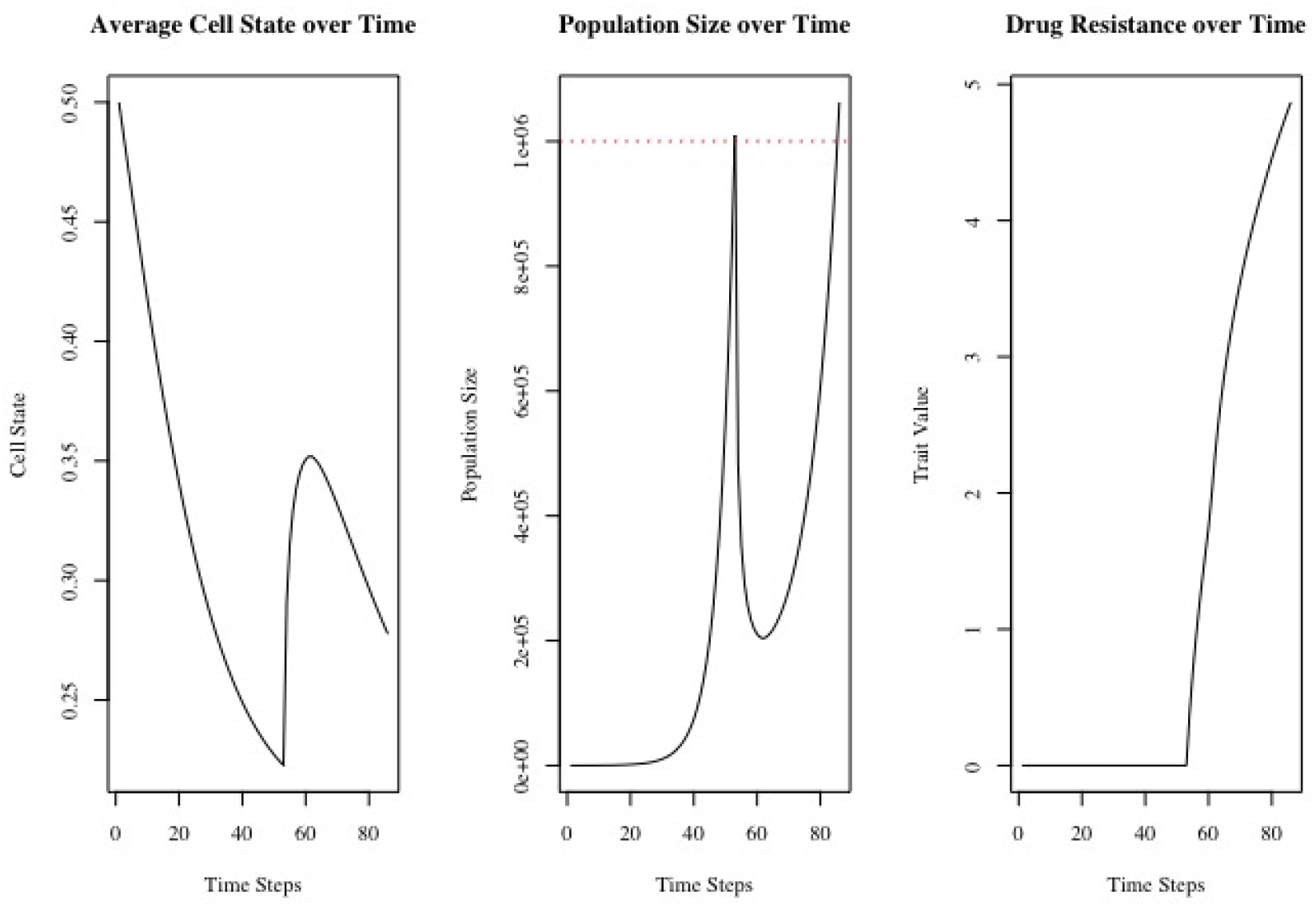
Impact of high evolvability (*k* = 4) on eco-evo-demo dynamics. Highly evolvable populations suffer a less drastic drop in population size upon the application of therapy and lead to shorter times to progression and treatment failure.

Once therapy is given, populations quickly shift their demography towards high cell states (Supplementary Movies 3,4). The high plasticity variance population is able to do this most rapidly and reaches the highest average cell state. Due to its higher average cell state pre-therapy, the initial effect of therapy is less severe than for the baseline and low plasticity variance cases. As resistance evolves, the population shifts back to a low cell state-biased distribution. Since the high plasticity variance population occupies higher cell states, the selection pressure to evolve resistance is less than that of the baseline and low variance cases. Thus, the higher plasticity variance population evolved the least resistance, but was able to progress and reach treatment failure the quickest (Table 2).

Next, we examine the contribution of evolvability to the eco-evo-demo dynamics of the population. Since we already simulated dynamics under no evolution in the prior section, we focus on the high evolvability case here. Due to the cost of evolvability, it takes longer for the population to pass the one million cell threshold. Once therapy is given, the population shifts its demography towards higher cell states. But due to the population’s high evolvability, it simultaneously evolves resistance quite rapidly. As a result, the state transition is not as dramatic as before, with the average cell state in the population reaching a maximum of only *u ≈* 0.35 before shifting back towards lower cell states (Supplementary Movie 5). As one might expect, the rapid evolution of resistance leads to a higher minimum of the population and a shorter TTP and TTF than other cases (Table 2).

### 3.3 Facultative Stress-Induced Responses

With a basic understanding of how variance and evolvability impact eco-evo-demo dynamics, we now examine facultative stress-induced responses that populations may undergo. Throughout this paper, we have discussed two mechanisms by which cells can adapt to a therapeutic stressor: by changing their cell state (plasticity) and by evolving higher resistance levels (genetic evolution). However, both of these adaptations come with a cost. A shift in demography towards higher cell states leads to lower proliferation rates, and a faster rate of evolution (via a high evolvability) leads to a higher death rate. Thus, it is natural to ask: What if populations could modulate their degree of plasticity and evolvability depending on their condition? In other words, what if cells could increase their plasticity and evolvability when they are stressed and decrease them when they are stable? In this section, we consider how facultative plasticity and adaptive mutagenesis impact the eco-evo-demo dynamics of populations under therapy.

Plasticity, the process by which cells transiently adopt a distinct phenotypic identity in a non-genetic fashion, is a commonly observed stress response in cancer and bacteria. Critically, these plastic transitions are completely reversible. In bacteria, a slow-cycling drug-tolerant persister (DTP) state plays such a role, providing a steppingstone towards more permanent genetic resistance [89]. A similar DTP state was detected in non-small cell lung cancer, melanoma, and glioblastoma [6, 60, 75, 76, 85, 91]. In addition, the ADRN-MES transition in neuroblastoma, epithelial-to-mesenchymal transition, and polyaneuploid transition (PAT) have all been shown to contribute to therapeutic resistance in cancer [3, 18, 55, 59, 71, 72, 101, 107, 109].

To implement facultative plasticity, we use the same parameter values as in Table 1, but let the variance in the plasticity kernel depend on the cell’s condition: 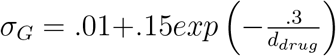 where 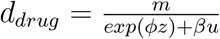 represents the probability of death for a cell in state *z* with resistance level *u* due to drug treatment. In addition to the ecoevo-demo dynamics, we plot the change in average plasticity variance over time (Fig. 5).

**Figure 5:**
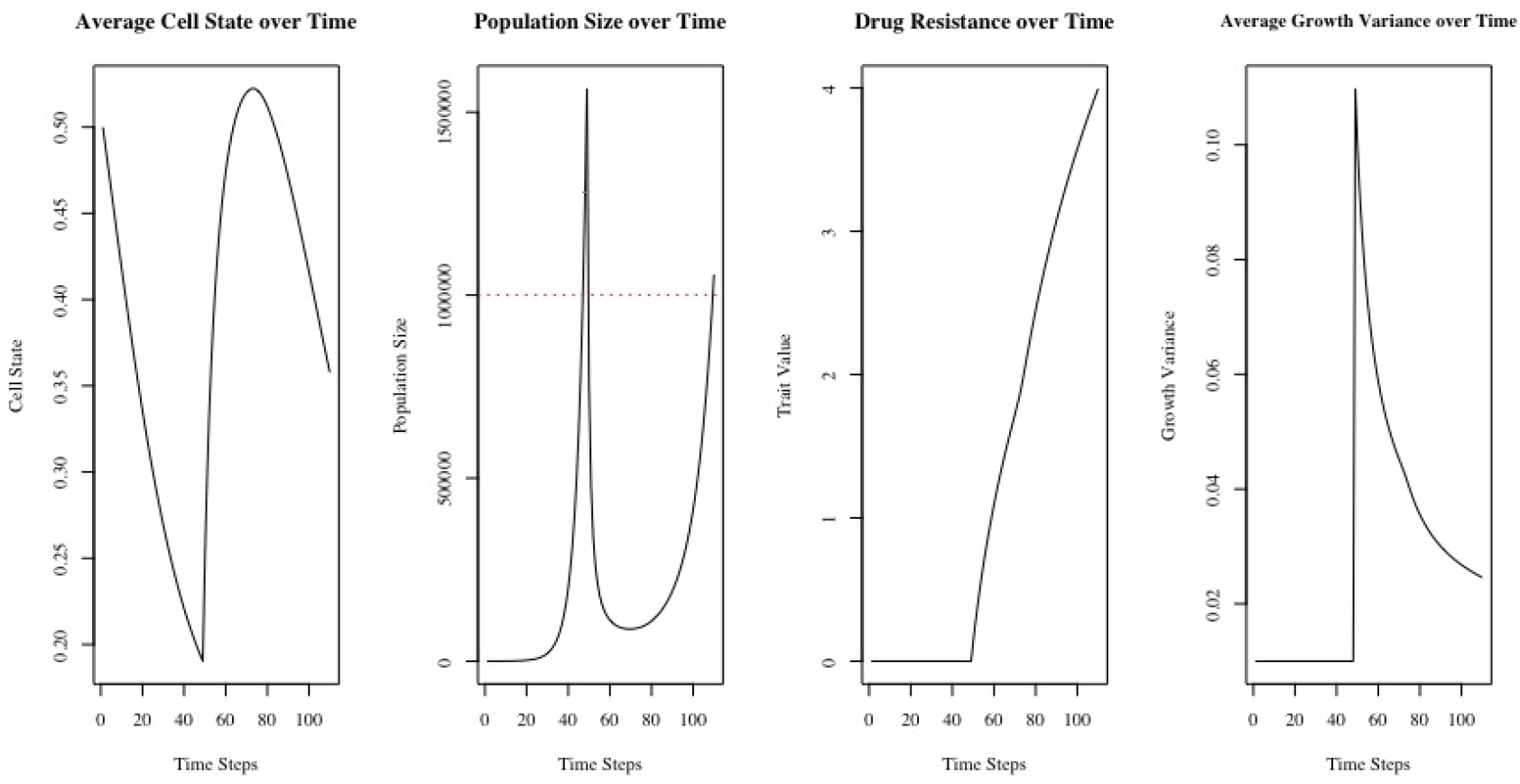
Impact of facultative plasticity on eco-evo-demo dynamics. The ability of a population to modulate the extent of its plasticity in response to its condition leads to faster proliferation pre-therapy, higher minima, and shorter times to progression and treatment failure.

Before drug exposure, the variance in plasticity kernel is low. This allows the population to reach lower average cell states and proliferate more quickly. Once therapy is applied, the population is able to rapidly increase its plasticity variance and move towards higher cell states (Supplementary Movie 7). Due to this shift in demography and increase in variance (that promotes a rapid evolution of resistance initially), the population drop is less severe (Table 3). Once the initial burst of adaptation and evolution has occurred, the population decreases its plasticity variance, allowing for continued modest evolution of resistance, and shifts its demography back towards lower states, allowing for more rapid proliferation. These aspects contribute to a faster rate of evolution and a shorter TTP and TTF compared to the baseline, low variance, and high variance cases.

**Table 3:** Facultative plasticity and stress-induced mutagenesis results summary.

|  | Facultative Plasticity | Adaptive Mutagenesis | Directed Mutagenesis |
| --- | --- | --- | --- |
| Drug Resistance | 4.000 | 3.992 | 3.469 |
| Minimum of Population | 87954 | 283292 | 395006 |
| Start of Therapy | 48 | 46 | 46 |
| Time to Progression | 62 | 31 | 23 |
| Time to Treatment Failure | 23 | 11 | 9 |

Next, we investigate adaptive mutagenesis, a widely recognized phenomenon in bacterial [7, 34, 32, 82], yeast [38, 49, 56, 61, 62, 70], and (albeit to a lesser extent) cancer [46, 30, 81, 93] evolution. Also known as stress-induced mutations, adaptive mutations describe the generation of heritable variation that occurs in response to the environment [80]. To implement adaptive mutagenesis, we use the same parameters in Table 1, but let evolvability be a function of the cell’s condition: 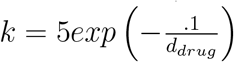 (Fig. 6A). Another related but controversial idea is that of directed mutagenesis in which useful mutations are induced preferentially (or at least that mutations occur more frequently in genes that improve fitness if mutated) [13, 88]. This is implemented in the same way as adaptive mutagenesis, except we remove a cost of evolvability altogether (Fig. 6B).

**Figure 6:**
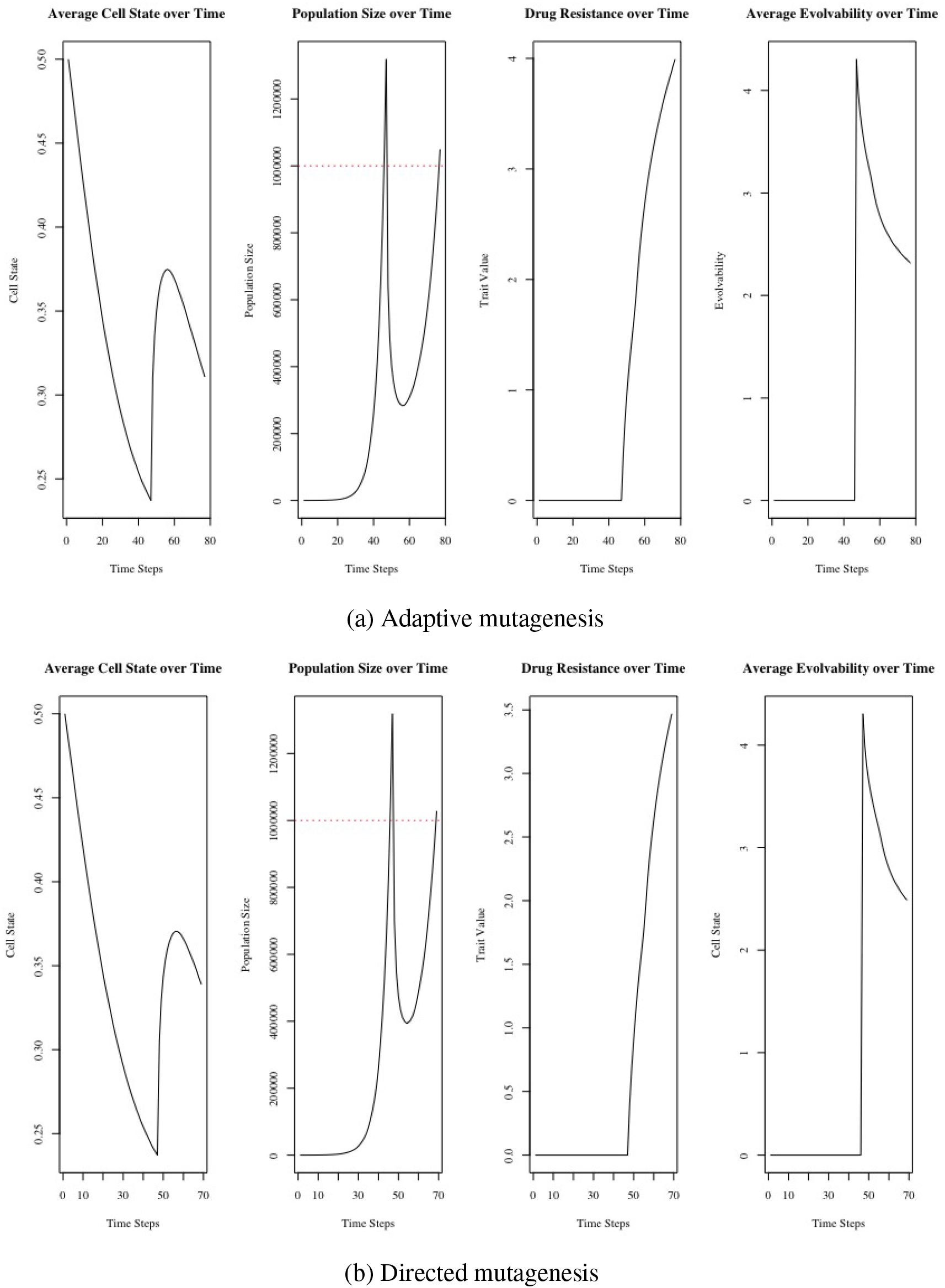
Impact of adaptive mutations on eco-evo-demo dynamics. The ability of populations to modify the nature of their mutations depending on their condition allows them to survive more effectively pre-therapy and evolve rapidly to the therapeutic stressor, leading to high minima and short times to progression and treatment failure.

Although a simplistic formulation, since evolvability is set to zero during the pre-therapy period, the cells do not suffer a natural death rate and their dynamics are identical to no evolution scenario. Once therapy is given, the population increases its evolvability and rapidly evolves resistance. Since evolvability is allowed to exceed *k* = 4 for a short period of time, the population experiences a higher minimum and a shorter TTP and TTF than the high evolvability case (Table 3). As resistance is acquired, the population gradually decreases its evolvability due to the trade-off between death due to drug treatment and natural death rate. The directed mutagenesis case is qualitatively identical to the adaptive mutagenesis case, but is quantitatively more exaggerated since the natural death rate is zero, regardless of evolvability.

### 3.4 Therapeutic Strategies

Throughout this paper, we have seen how the potent combination of plasticity and genetic evolution allows populations to evolve and adapt to stressful conditions. This leads to the natural question: How can we develop therapeutic strategies to effectively control or drive populations to extinction given the complex interplay among ecological, evolutionary, and demographic dynamics? As done in our past work and as suggested by others, one approach is to use a life-history enlightened approach [11, 12, 90] by attempting to block plastic transitions towards drug-resistant phenotypes. Such approaches, at the epigenetic, intracellular pathway, and microenvironmental levels, have shown promise in a variety of cancers, including glioblastoma, breast cancer, melanoma, and prostate cancer [45, 60, 83, 84, 85, 94, 100]. Similarly, efforts have been made to develop drugs to target cell states that are resistant to traditional therapies [8, 48, 76, 96, 99]. In this section, we focus on therapeutic strategies assuming that neither of these options exist.

Specifically, we will investigate the efficacy of intermittent and adaptive therapy protocols when cells can adapt to the stressor in both plastic and genetic ways. For simplicity, we will implement therapy solely on the baseline case (see Table 1) here. Under both protocols, we let therapy begin once the population size exceeds one million cells. We focus on three statistics: minimum of population, TTF, and TTP. If extinction is the goal, the minimum of the population is the metric of most importance. If the physician is aiming at control and only has one drug, TTP is most relevant. However, if the goal is control several drugs are available, TTF is key.

First, we begin with the intermittent therapy. Under this protocol, therapy follows a fixed administration schedule, with both treatment and drug holiday times set constant for the entirety of the simulation. Once again, we set the drug dosage to be constant for the entirety of the treatment periods. Using the parameter values from Table 2, we simulate a sample intermittent protocol with 8 time steps on therapy and 4 time steps off, arbitrarily chosen (Fig. 7). The effects of the intermittent nature of the protocol can clearly be seen reflected in the ecological, evolutionary, and demographic dynamics of the population (Supplementary Movie 9). Before progression, during periods of therapy, the population size decreases, resistance evolves, and the population shifts towards lower cell states. As cells gain resistance however, each of these trends becomes less severe. Once progression occurs, the population is able to proliferate and reduce their cell state even in the presence of therapy (albeit in a less effective manner).

**Figure 7:**
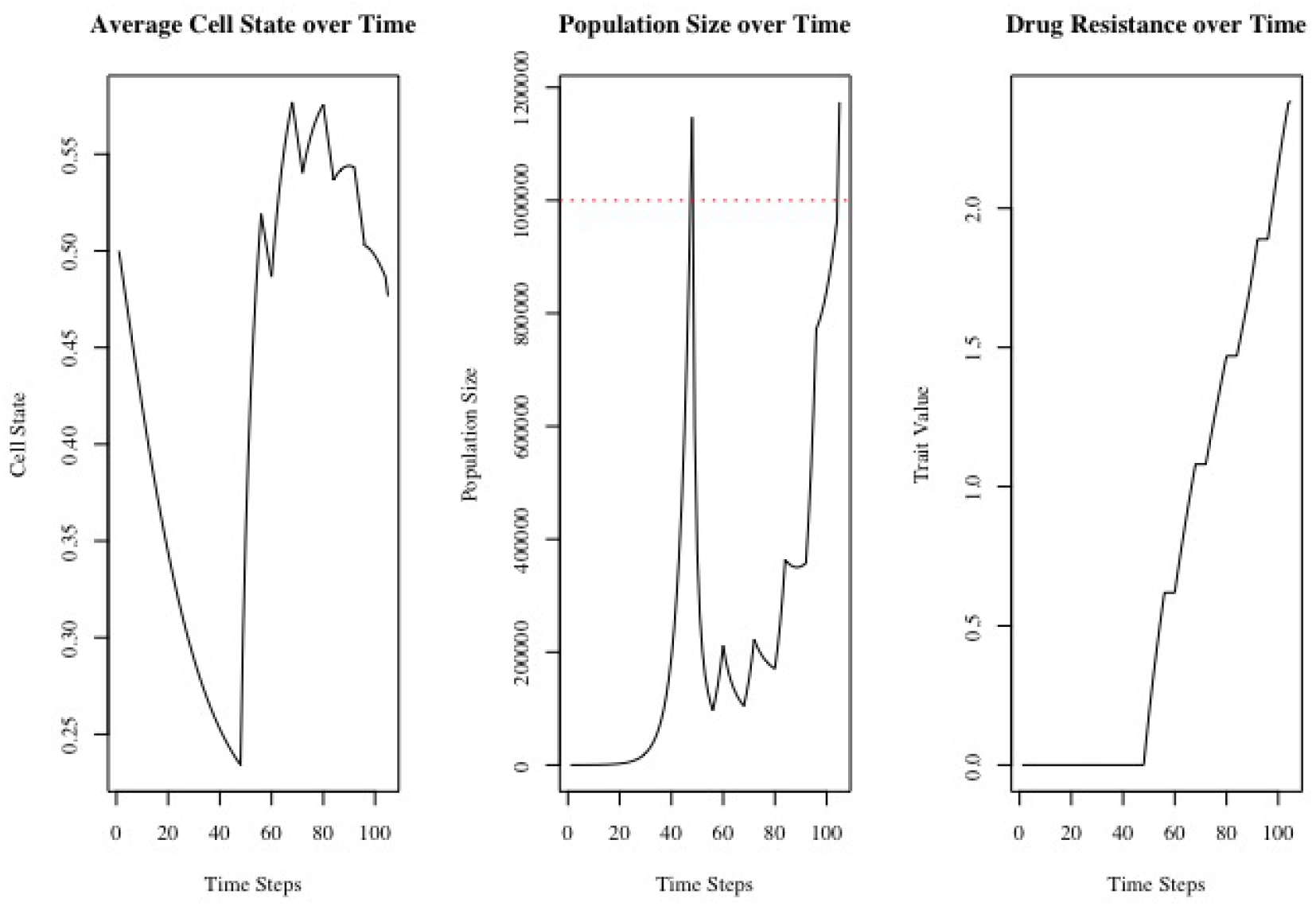
Intermittent therapy protocol. Longer holidays/shorter treatment times lead to higher minima, a shorter time to progression, but a longer time to treatment failure.

To better understand the effects of intermittent therapy, we simulate a range of treatment times and drug holidays and summarize key statistics in Fig. 8. Greater drug holiday durations and lower treatment times lead to higher minima as the therapy is not given enough time to reduce the population down to its minimum. Longer holidays and shorter treatment durations also lead to shorter TTP (since the population recovers to higher levels) and a longer TTF (since resistance evolves less quickly). Thus, assuming no cost of resistance, continuous therapy is more effective for extinction and control purposes when only one drug is available. On the other hand, including drug holidays may be more effective for multi-drug control.

**Figure 8:**
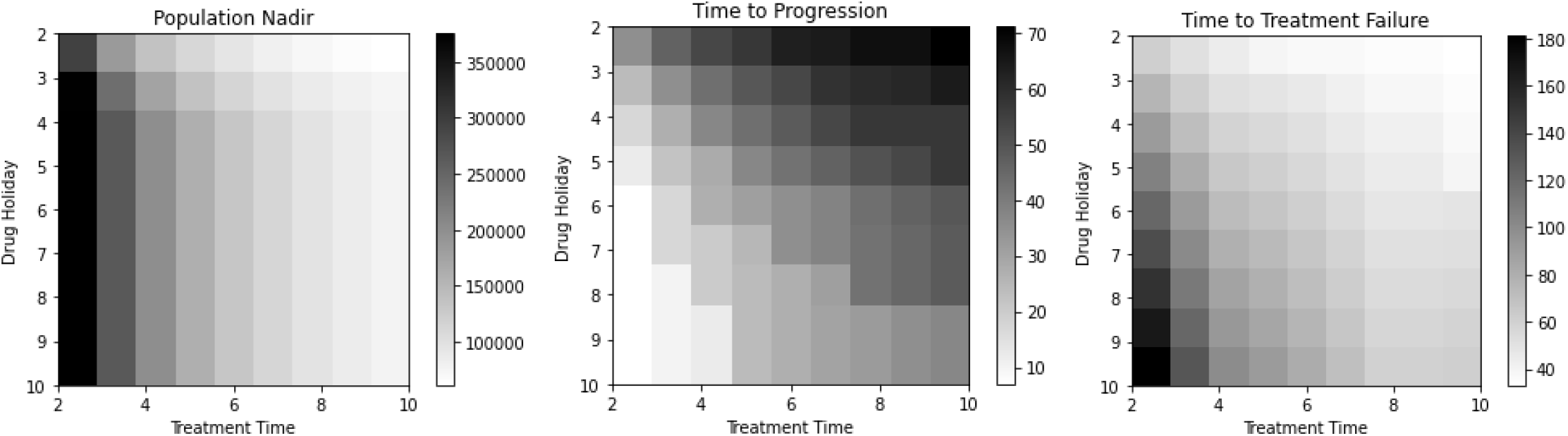
Intermittent therapy: minimum of population, time to progression, and time to failure.

Next, we turn to our adaptive therapy protocol. Under this protocol, we administer therapy when the population increases past a threshold and remove therapy when the population decreases below another threshold. We first create a sample adaptive therapy protocol by setting the upper threshold to 800,000 cells and the lower thresh-old to 400,000 cells, arbitrarily chosen (Fig. 9). As before, we can clearly see the impacts of treatment times and treatment holidays on the eco-evo-demo dynamics of the population (Supplementary Movie 10). Due to the adaptive nature of the dosing, we note that switching treatment on and off is more frequent at the beginning of therapy, when cells are relatively more sensitive to the drug. As cells gain resistance, therapy must be given for longer periods of time to push the population below the lower threshold. As before, drug dosage is held constant during treatment periods.

**Figure 9:**
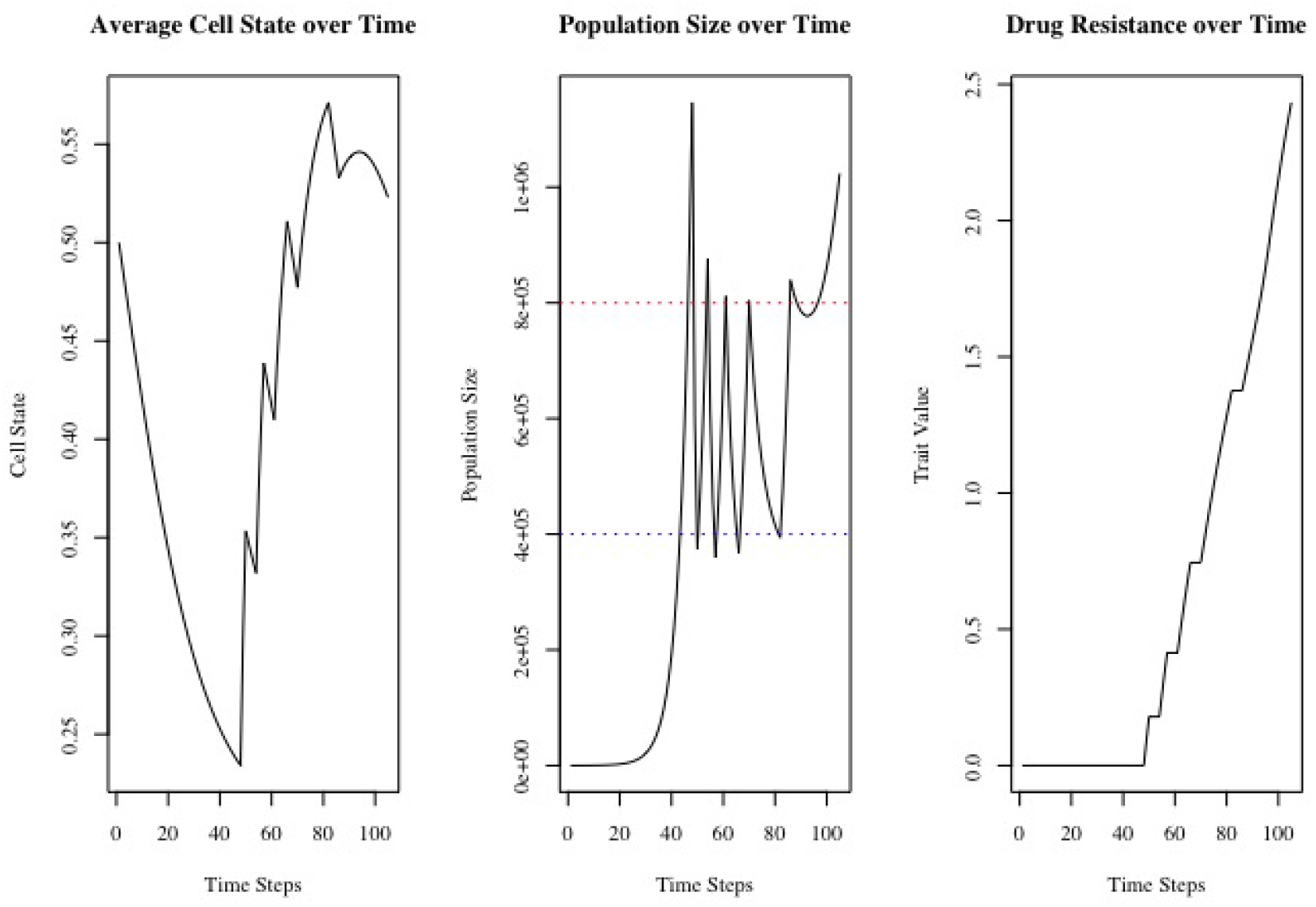
Adaptive therapy protocol. Higher thresholds for starting and stopping therapy leads to a shorter time to progression and a longer time to treatment failure.

To gain a deeper understanding on how the thresholds we set impact TTP and TTF, we simulate adaptive therapy protocols over a range of thresholds (Fig 10). As we can see, higher thresholds for starting and stopping therapy lead to a shorter TTP and a longer TTF. High thresholds induce more therapy cycles than lower thresholds because less therapy is required to reduce the population down to the threshold bounds. This prolongs the time required to gain resistance; however, since the population is cycling at high levels, progression is not far once the cells escape.

**Figure 10:**
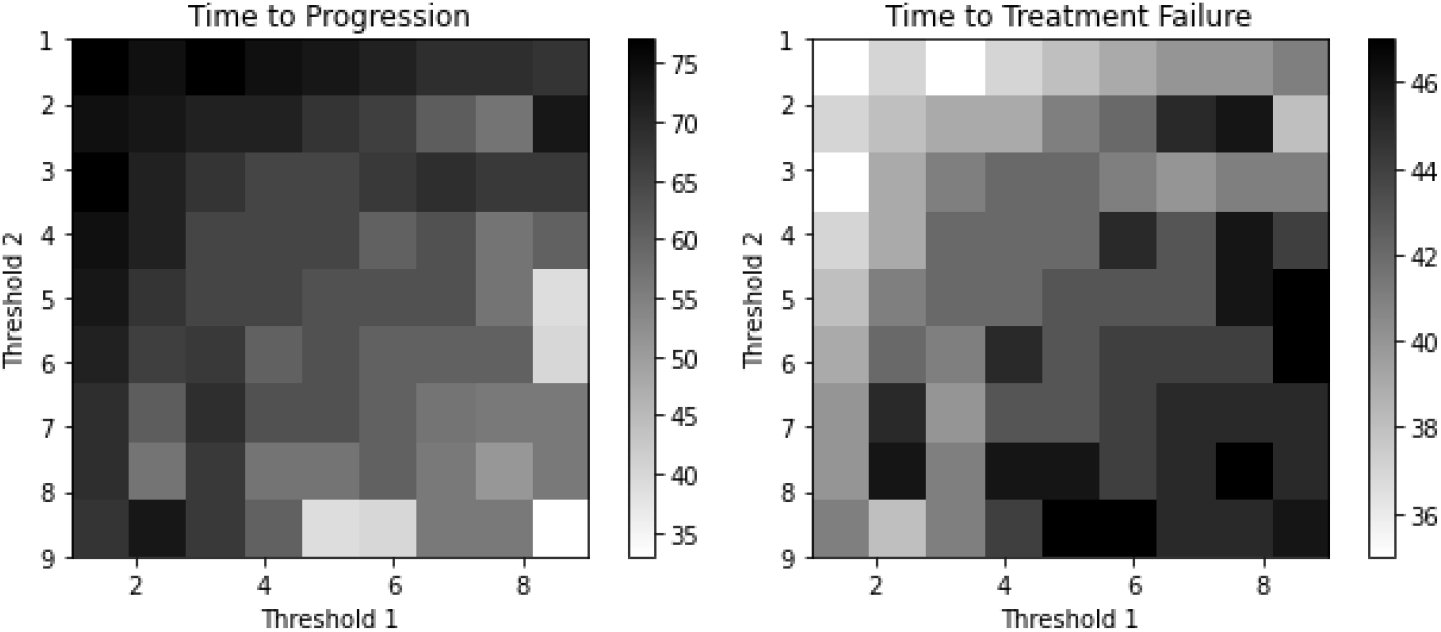
Impact of thresholds on adaptive therapy protocols: time to progression and time to failure.

Under this model formulation, we assume no cost of resistance. This leads to an infinite improvement model of evolution, in which *u* is monotonically increasing. While this is a valid assumption in some cases, many cancers exhibit such costs. If we instead include a cost of resistance (as exists in many cancers [39, 53, 54, 92]), we expect *u* to decrease during times of drug holidays and become re-sensitized, at least to some degree, to therapy upon the next administration. This could conceivably prolong TTP and TTF and may change our qualitative observations.

To test this hypothesis, we let *b*(*u,z*) = *b*_1_*exp*(*b*_2_*z* + *b*_3_*u*) where *b*_3_ = *−*0.2. This induces a proliferation-resistance trade-off. For biological realism, we force *u ≥* 0 to prevent negative resistance values pre-therapy. Using a drug dosage of *m* = 0.7, we simulate continuous therapy, intermittent therapy, and adaptive therapy protocols (Fig. 11). For intermittent therapy, therapy is on for 8 time steps, then turned off for 4 time steps. For adaptive therapy, we use an upper threshold of 800,000 cells and a lower bound of 400,000 cells.

**Figure 11:**
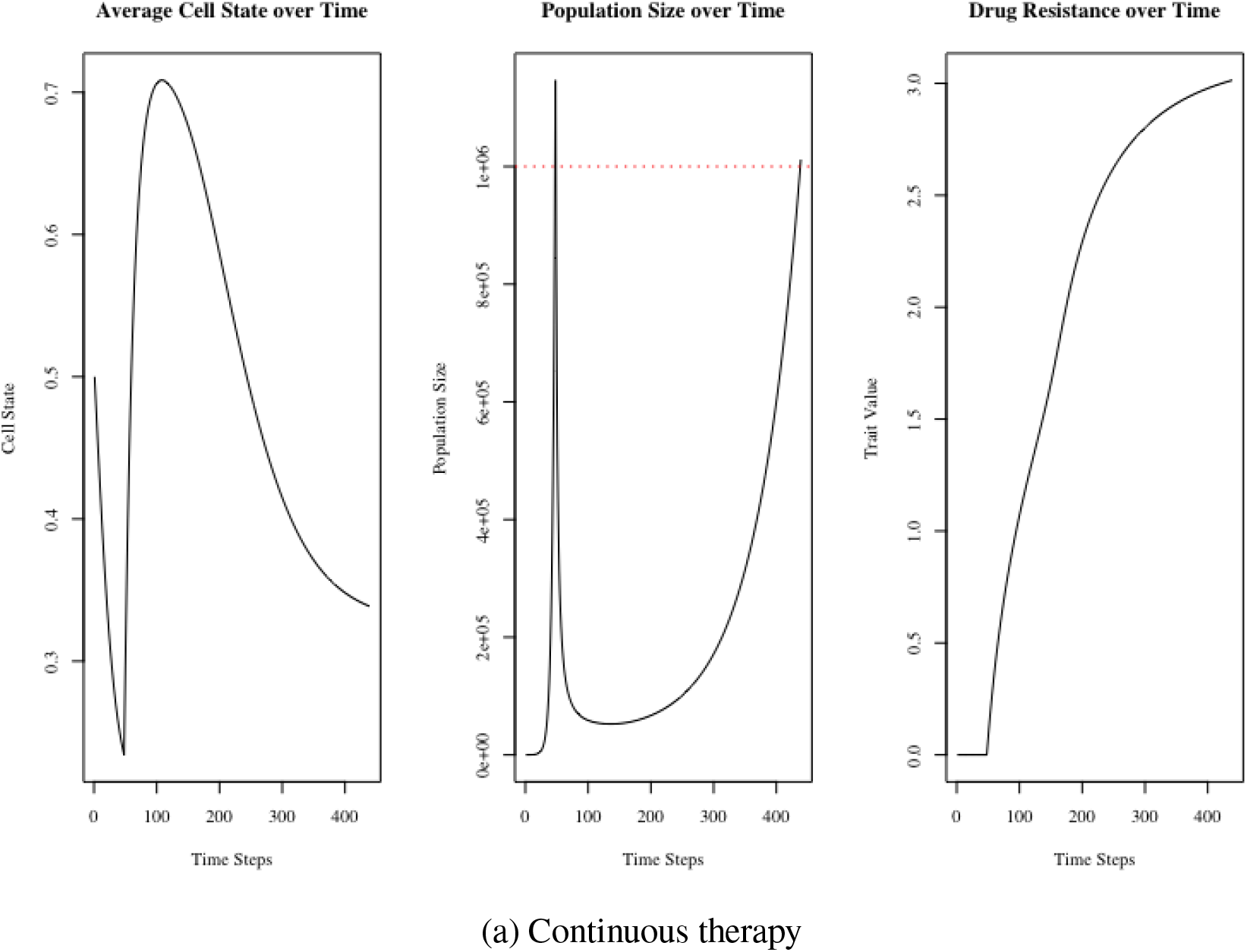
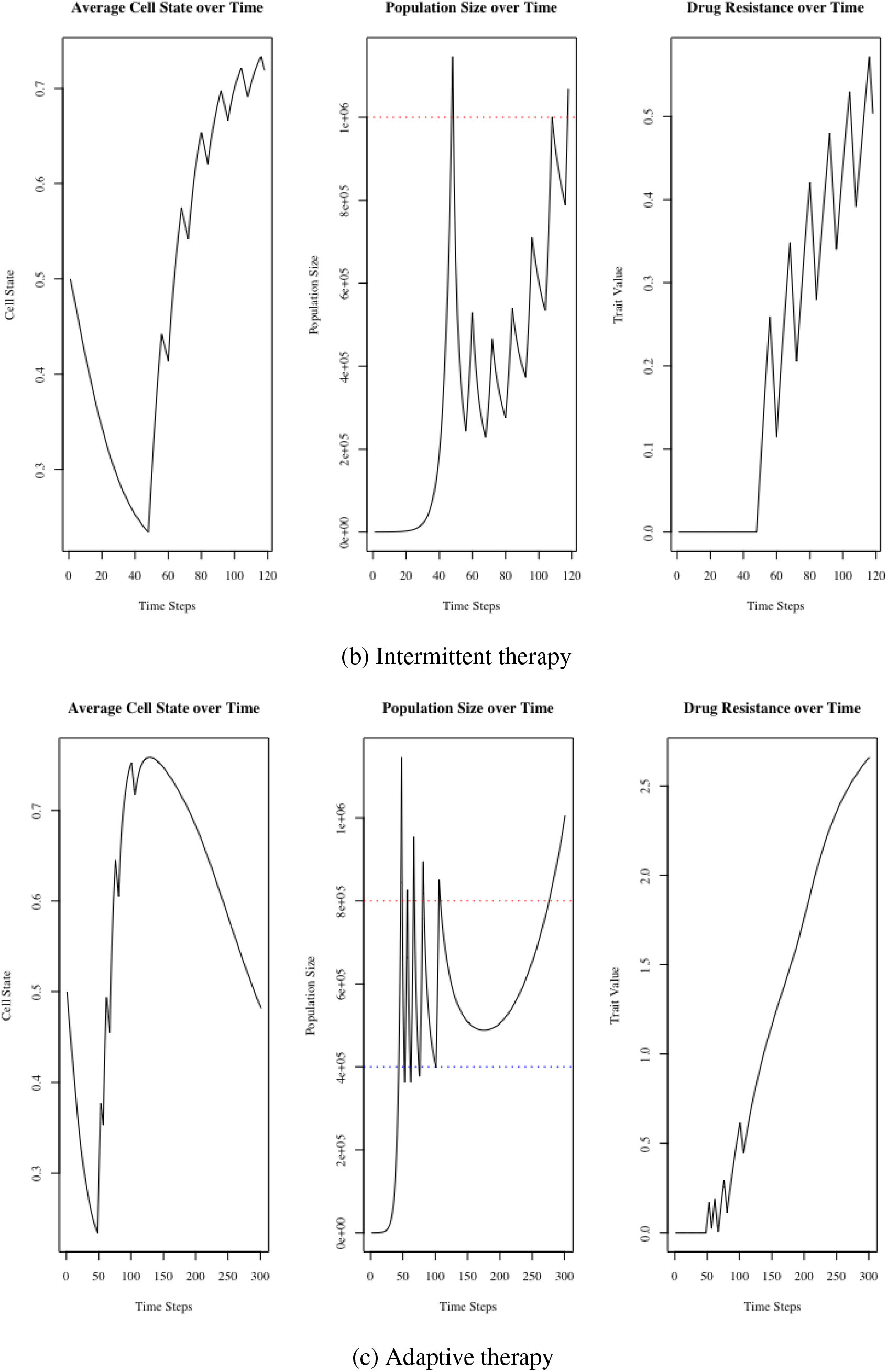
Impact of cost of resistance on continuous, intermittent, and adaptive therapy protocols. The cost of resistance leads to higher times to progression and failure and lower resistance levels across the board. However, the same qualitative trends hold, with continuous therapy leading to a lower population minimum and higher time to progression whereas intermittent and adaptive therapies lead to higher times to failure.

Due to the cost of resistance, during times of therapy, *u* increases, but during times of drug holiday, *u* decreases, leading to resensitization of therapy (Supplementary Videos 11-13). In all cases, this leads to lower overall levels of resistance and prolongs TTF and TTP (Table 4). However, even with this cost, we notice the same general trends as before: Continuous therapy is more effective at producing a low minimum of the population and promotes the longest TTP. On the other hand, intermittent and adaptive therapies are effective at keeping resistance levels low in the population, promoting a longer TTF.

**Table 4:** Continuous, intermittent, and adaptive therapy with cost of resistance summary.

|  | Continuous Therapy | Intermittent Therapy | Adaptive Therapy |
| --- | --- | --- | --- |
| Drug Resistance | 3.014 | 0.504 | 2.660 |
| Minimum of Population | 52585 | 228928 | 488300 |
| Time to Progression | 391 | 70 | 253 |

## 4 Discussion

In this paper, we developed a novel theoretical framework using IPMs to examine stress-induced responses, particularly in bacterial and cancer cell populations. This framework allowed us to simultaneously consider the roles of plasticity (in a continuous, rather than discrete sense) and gradual genetic evolution in a population’s ability to respond to stressful environments. We found that faster evolving (i.e., highly evolvable) and more plastic populations (i.e., those with a high plasticity variance) are able to respond to stress effectively, avoiding extinction and causing treatment failure and progression rapidly. We then branched into facultative stress-induced responses, in which plasticity and evolvability were condition-dependent. We found that these mechanisms were even more effective at promoting evolutionary rescue in the population and led to faster proliferation pre-therapy, higher minima, and shorter times to progression and treatment failure. We then considered intermittent and adaptive therapy regimens. Although continuous therapy is generally more effective if the goal is extinction or control with a single drug, introducing drug holidays can prolong TTF, an important aspect if the goal is multi-drug control.

Our study has several limitations. Being theoretical in nature, our parameter values were not informed by data. Future work could seek to calibrate and compare models with time-series RNA sequencing (at bulk or singlecell resolution) and population count data pre-therapy, during therapy, and after therapy. Furthermore, complementary methods such as agent-based modeling techniques could be employed for finer resolution and stochasticity. Different forms of the cost of evolvability and resistance, biased kernels for adaptive mutations and directed plastic transitions, and density dependence are all things future work can consider.

Theoretically, we hope to expand our modeling framework to include several continuous states. For instance, we could model the stochastic reversal of evolutionary resistance (a plausible outcome when a resistant population is left without therapy for extended periods of time) by treating resistance as a second state. We also hope to include hybrid discrete and continuous states, a notion particularly important when incorporating a polyploid state [12, 11] or considering primary and metastatic tumors or habitat heterogeneity within a tumor [21]. Finally, we hope that future work will apply this framework to the specific problems mentioned here: the epithelial-to-mesenchymal transition, ADRN-MES transition in neuroblastoma, and the PAT transition, as well as additional biological conundrums such as the stem-cell paradigm in wound healing and macrophage polarization.

## 5 Data Availability

Codes used to produce plots in this paper can be found at https://github.com/abukkuri/IPM_StressResponses.

## 7 Acknowledgements

The author would like to thank Stina Andersson, Joel S. Brown, and Sofie Mohlin for valuable conversations related to this project.

